# Profiling of oral microbiota and cytokines in COVID-19 patients

**DOI:** 10.1101/2020.12.13.422589

**Authors:** Valerio Iebba, Nunzia Zanotta, Giuseppina Campisciano, Verena Zerbato, Stefano Di Bella, Carolina Cason, Sara Morassut, Roberto Luzzati, Marco Confalonieri, Anna Teresa Palamara, Manola Comar

## Abstract

SARS-CoV-2 presence has been recently demonstrated in the sputum or saliva, suggesting how the shedding of viral RNA outlasts the end of symptoms. Recent data from transcriptome analysis show that oral cavity mucosa harbors high levels of ACE2 and TMPRSS2, highlighting its role as a double-edged sword for SARS-CoV-2 body entrance or interpersonal transmission. In the present study, for the first time, we demonstrate the oral microbiota structure and inflammatory profile of COVID-19 patients. Hospitalized COVID-19 patients and matched healthy controls underwent naso/oral-pharyngeal and oral swabs. Microbiota structure was analyzed by 16S rRNA V2 automated targeted sequencing, while oral and sera concentrations of 27 cytokines were assessed using magnetic bead-based multiplex immunoassays. A significant diminution in species richness was observed in COVID-19 patients, along with a marked difference in beta-diversity. Species such as *Prevotella salivae* and *Veillonella infantium* were distinctive for COVID-19 patients, while *Neisseria perflava* and *Granulicatella elegans* were predominant in controls. Interestingly, these two groups of oral species oppositely clustered within the bacterial network, defining two distinct Species Interacting Group (SIGs). Pro-inflammatory cytokines were distinctive for COVID-19 in both oral and serum samples, and we found a specific bacterial consortium able to counteract them, following a novel index called C4 firstly proposed here. We even introduced a new parameter, named CytoCOV, able to predict COVID-19 susceptibility for an unknown subject at 71% of power with an AUC equal to 0.995. This pilot study evidenced a distinctive oral microbiota composition in COVID-19 subjects, with a definite structural network in relation to secreted cytokines. Our results would pave the way for a theranostic approach in fighting COVID-19, trying to enlighten the intimate relationship among microbiota and SARS-CoV-2 infection.

## INTRODUCTION

Coronavirus disease 2019 (COVID-19) is a global pandemic established at the end of 2019, whose etiological agent is SARS-CoV-2, a member of betacoronaviruses ^1^. COVID-19 is typically characterized by i) symptoms of the lower respiratory tract ^2^; ii) a systemic ‘cytokine storm’ ^3^,4; iii) neurological symptoms such as ageusia and hyposmia ^2,5^. Patients exhibiting an exaggerated form of symptoms ^6^ showed greater levels of proinflammatory factors ^1^,7,8, and beside respiratory illnesses, they may also have enteric symptoms and encephalitis^5^. Virus replication in the throat particularly during the first 5 days of symptoms is strongly supported by identification of transcribed subgenomic mRNA in throat swab samples. However, some reports suggest the potential for pre- or oligosymptomatic transmission as consequence of a mild illness of the upper respiratory tract. In this context, understanding the invasive process of the virus at the port of entry represents the crucial point. The presence of SARS-CoV-2 have been recently demonstrated in the sputum or “posterior oropharyngeal saliva” ^9–12^ indicating the shedding of viral RNA outlasted the end of symptoms and the oral mucosa supports. Transcriptome analysis supports that expression of COVID-19 ACE2 and TMPRSS2 receptors in salivary glands and epithelial cells showing the potential vulnerability risk for oral cavity for lung or gut involvement ^13^. From these data, virus– host interplay within the oral cavity seems to be a promising feature of COVID-19 pathogenesis, forming the basis of disease severity and spread. The relationship between virus and host environment included disturbance of resident bacterial community, event recently reported also in gut from COVID-19 patients ^14^. As a consequence, a cascade of inflammatory markers is detected specifically as a local cytokines pathway. The characterization of such a relationship might identify physiological markers for the potential risk in terms of disease severity and therapeutic strategies. In the present study, for the first time, we characterized the interplay of COVID-19 infection with oral microbiota and inflammatory profile.

## MATERIALS AND METHODS

### Study Cohort and Samples

A total of 26 patients, 6 women (mean age 66±16 years) and 20 men (mean age 66±15 years) hospitalized at the Infectious Diseases Unit, University of Trieste, Italy, between April 10^th^ 2020 and May 5^th^ 2020, tested positive for COVID-19, were selected for this study. All patients provided informed consent for the use of their data and clinical samples for the purposes of the present study. Patients acquired their infections upon known close contact to an index case, thereby avoiding representational biases owing to symptom-based case definitions. All patients had interstitial pneumonia and were receiving oxygen therapy but did not require endotracheal intubation and invasive mechanical ventilation. Oropharyngeal and nasopharyngeal swabs for diagnosis of SARS-CoV-2 were performed, and oral swab specimens touching tongue, palate and cheeks were additionally collected two days after hospital admission for oral microbiota and local immune response characterization. No mouth washing products were administered to the patients. Specimens were additionally collected with the same modality from age-matched healthy volunteers (n=15) without evaluable risk for SARS-CoV-2 infection. In addition, sera samples from 11 infected patients with severe disease which underwent endotracheal intubation and invasive mechanical ventilation, were additionally analyzed for peripheral cytokines profile.

### SARS-CoV-2 Detection

SARS-CoV-2 detection was performed on the CFX96™ Real-Time PCR Detection System (Bio-Rad, California, USA), using the NeoPlex™ COVID-19 Detection Kit (Genematrix, Seongnam, Kyonggi-do, South Korea) targeting the viral N and RdRp genes and the housekeeping gene of β-actin as internal control, following the manufacturer’s instructions.

### Soluble Immune Mediators Quantification

The profile of a panel of 27 cytokines including chemokines and growth factors was assessed in duplicate, in oral swabs of positive and negative subjects for SARS-CoV-2 using magnetic bead-based multiplex immunoassays (Bio-Plex Pro™ human cytokine 27-plex panel, Bio-Rad Laboratories, Milan, Italy) according to the pre-optimized protocol ^15^. Briefly, the undiluted samples (50 μl) were mixed with biomagnetic beads in 96-well flat-bottom plates, and after incubation for 30 min at room temperature followed by washing plate with Bio-Plex wash buffer, 25 μl of the antibody–biotin reporter was added. After the addition of 50 μl of streptavidin– phycoerythrin (PE) and following incubation for 10 min, the concentrations of the cytokines were determined using the Bio-Plex-200 system (Bio-Rad Corp., United States) and Bio-Plex Manager software (v.6, Bio-Rad). The data were expressed as median fluorescence intensity (MFI) and concentration (pg/ml).

### ACE2 and TMPRSS2 expression

The expression levels of human ACE2 and TMPRSS2 genes were evaluated by SYBR green PCR analyses. In brief, RNA was reverse transcribed using SensiFast cDNA Synthesis Kit (Bioline), and SYBR green PCR analysis was performed using Kapa HiFi HotStart Ready Mix (Roche) ^16^. The housekeeping Beta-globin human gene was used for normalization and the relative expression levels (ΔCt) of human ACE2 and TMPRSS2 genes were compared between groups.

### DNA and RNA extraction

Total DNA and RNA were extracted starting from 300 μl and 200 μl of samples respectively, in a final elution volume of 50 μl, using the Maxwell CSC Instrument (Promega Srl, Italy) and following the manufacturer’s instructions. 50 μl of oral samples were additionally aliquoted for profiling and quantification of soluble cytokines.

### 16S targeted sequencing

The V2–V3 portion of the 16S rRNA was amplified, using the primer set F101-R534,with a different IonXpress barcode per sample attached to the reverse primer. PCR reactions were performed using the Kapa Library Amplification Kit (Kapa Biosystems, Massachusetts, USA) and BSA 400ng/μL, under the following conditions: 5 min at 95°C, 30 sec at 95°C, 30 sec at 59°C, 45 sec at 72°C and a final elongation step at 72°C for 10 min. DNA after normalization was quantified with a Qubit^®^ 2.0 Fluorometer (Invitrogen, Carlsbad, California, USA). The amplicon size was checked on a 2% agarose gel. The subsequent step of PCR purification was carried out using the Mag-Bind^®^ Total Pure NGS beads (OMEGA Bio-Tek, Georgia, USA), retaining fragments >100 bp. Template preparation was performed by the Ion PGM Hi-Q View kit on the Ion OneTouch™ 2 System (Life Technologies, Grand Island, NY, United States) and sequenced using the Ion PGM Hi-Q View sequencing kit (Life Technologies, Grand Island, NY, United States) with the Ion PGM™ System technology. Negative controls, including a no-template control, were processed with the clinical samples ^17^.

### Microbiota characterization

Raw FASTQ files were analyzed with DADA2 pipeline v.1.14 for quality check and filtering (sequencing errors, denoising, chimera detection) on a Workstation Fujitsu Celsius R940 (Fujitsu, Tokyo, Japan) (Figure S1). Filtering parameters were as follows: truncLen=0, minLen=100, maxN=0, maxEE=2, truncQ=11, trimLeft=15. All the other parameters in the DADA2 pipeline for single-end IonTorrent were left as default. Raw reads (2447325 in total, on average 59691 per sample) were filtered (818531 in total, on average 19964 per sample) and 962 Amplicon Sequence Variants (ASV) were found. Sample coverage was computed and resulted to be on average higher than 99% for all samples, thus meaning a suitable normalization procedure for subsequent analyses. Bioinformatic and statistical analyses on recognized ASV were performed with Python v.3.8.2. Each ASV sequence underwent a nucleotide Blast using the National Center for Biotechnology Information (NCBI) Blast software (ncbi-blast-2.3.0) and the latest NCBI 16S Microbial Database accessed at the end of July 2020 (ftp://ftp.ncbi.nlm.nih.gov/blast/db/). After blasting, the 962 ASVs were merged into 122 species (thus excluding sub-species or strain differences), and a matrix of their relative abundances was built for subsequent statistical analyses.

### Network analysis

Pearson matrices for network analysis (metric = Bray-Curtis, method = complete linkage) were generated on normalized and standardized data with in-house scripts (Python v3.8.2) and visualized with Gephi v.0.9.2, as previously reported ^18^. Bacterial species having a prevalence ≥5% were considered to generate the nodes within the final network, while a significant Pearson correlation coefficient and its related *P* value (after Benjamini-Hochberg FDR at 10%) was employed to obtain eight categories defining edge thickness ^19^. A leave-one-out method was employed by SciKit-learn package v0.4.1 on the subjects in order to have an averaged *P* value for each correlation among two definite variables. Network analysis was performed on unified datasets ^18^ taking care of an optimized visual representation with Gephi v.0.9.2, as proposed by current guidelines ^20–24^. Nodes were colored according to the cohort in which species harbored the highest mean relative abundance, after normalization and standardization. The degree value, measuring the in/out number of edges linked to a node, and the betweenness centrality, measuring how often a node appears on the shortest paths between pairs of nodes in a network, were computed with Gephi v.0.9.2. Intranetwork communities (here called Species Interacting Groups - “SIGs” ^18^,25) were retrieved using the Blondel community detection algorithm ^26^ by means of randomized composition and edge weights, with a resolution equal to 1 ^27^.

### Statistical analysis

Data matrices (microbiota taxa or cytokines) were firstly normalized then standardized using QuantileTransformer and StandardScaler methods from Sci-Kit learn package v0.20.3. Normalization using the output_distribution=‘normal’ option transforms each variable to a strictly Gaussian-shaped distribution, whilst the standardization results in each normalized variable having a mean of zero and variance of one. These two steps of normalization followed by standardization ensure the proper comparison of variables with different dynamic ranges, such as bacterial relative abundances, or cytokines levels. For microbiota analysis, measurements of α diversity (within sample diversity) such as Richness and Shannon index, were calculated at species level using the SciKit-learn package v.0.4.1. Exploratory analysis of β-diversity (between sample diversity) was calculated using the Bray-Curtis measure of dissimilarity and represented in Principal Coordinate Analyses (PCoA), along with methods to compare groups of multivariate sample units (analysis of similarities - ANOSIM, permutational multivariate analysis of variance - PERMANOVA) to assess significance in data points clustering ^28^. ANOSIM and PERMANOVA were automatically calculated after 999 permutations, as implemented in SciKit-learn package v0.4.1. We implemented Partial Least Square Discriminant Analysis (PLS-DA) and the subsequent Variable Importance Plot (VIP) as a supervised analysis wherein the VIP values (order of magnitude) are used to identify the most discriminant bacterial species among COVID-19 and control samples. Bar thickness reports the fold ratio (FR) value of the mean relative abundances for each species among the two cohorts, while an absent border indicates mean relative abundance of zero in the compared cohort. In order to compare the microbiota species with cytokines levels, a multivariate statistical Pearson correlation analysis (and related P values) was performed with custom scripts (Python v3.8.2), and a Hierarchical Clustering Analysis (HCA) with ‘Bray-Curtis’ metrics and ‘complete linkage’ method was used to visualize putative cross-correlation clusters. Mann-Whitney U and Kruskal-Wallis tests were employed to assess significance for pairwise or multiple comparisons, respectively, considering a P value <0.05 as significant. Statistical analyses gathering more than two groups were performed using ANOVA followed by pairwise comparisons with Bonferroni adjustments. Differential enrichment analyses in murine studies were corrected for multiple hypothesis testing using a two-stage Benjamini-Hochberg FDR at 10%.

## RESULTS

Data relative to demographics and clinical data of enrolled patients at the time of samples collection were resumed in Supplementary Table 1. Cardiac dysfunctions (14/26) and neurological involvement (11/26) including ageusia or hyposmia (9/26) and paralysis and epilepsy (2/26) represent the most frequently present comorbidities. Regarding drug therapies, all patients were treated with hydroxychloroquine and 57.7% of them (15/26) received combinations with antibiotics.

### COVID19 patients harbor a distinctive oral microbiota

Following 16S targeted sequencing, we observed a significant diminution (−40%) of alpha-diversity (species richness) in COVID-19 patients (*P*=2.92*10^−2^) (Figure 1A), while Shannon biodiversity was unaltered. The unsupervised algorithm of Principal Coordinate Analysis (PCoA) visually represented a significant separation of COVID-19 oral samples from controls (P<1*10^−3^) (Figure 1B), thus meaning a different oral microbiota composition assessed with two different measures of beta-diversity (ANOSIM and PERMANOVA). In order to find a pattern of bacterial species able to describe the changes in microbiota composition of COVID-19 samples, we used the supervised algorithm of Partial Least Squares - Discriminant Analysis (PLS-DA), which generated a Variable Importance Plot (VIP) showing the most important species able to separate the two cohorts (Figure 1C). Six bacterial species, having a VIP score higher than the chosen cutoff of 1.25, were discriminant for COVID-19 (*Haemophilus parainfluenzae*, *Veillonella infantium*, *Soonwooa purpurea*, *Prevotella salivae*, *Prevotella jejuni*, *Capnocytophaga gingivalis*), while several species (n=23) were significantly distinctive for controls (the most important being *Neisseria perflava*, *Lampropedia puyangensis*, *Rothia mucilaginosa*, *Kallipyga gabonensis*, *Candidatus Flaviluna*, *Granulicatella elegans*). Interestingly, through network analysis we were able to retrieve four communities, also known as Species Interacting Groups (SIGs) (Figure 1D), which represent a topological clusterization of bacterial species linked to “functional modules”, for example a disease status. In order to see if SIGs would be related to COVID-19, nodes were colored according to the cohort in which species had the highest mean relative abundance (after normalization and standardization) (Figure 1D). Two SIGs (SIG1, SIG4) harbored the majority of COVID-19-related species (18/22, 82%), while SIG2 and SIG3 contained mostly controls species (19/20, 95%), and this repartition of species was significant (Fisher test with Freeman-Halton extension, *P*=2.13*10^−6^). SIG1 harbored three out of six COVID-19-related species extrapolated from VIP plot (namely, *Veillonella infantium*, *Prevotella salivae*, *Prevotella jejuni*), while *Soonwooa purpurea* was included in SIG4, along with other two species known for their noxious effects (*Atopobium parvulum* and *Fusobacterium nucleatum*). The first species depicted by VIP plot as discriminant for COVID-19, namely *Haemophilus parainfluenzae*, was indeed colored as control within the network because of the normalization/standardization procedure, thus meaning that this species would be not reliable as a descriptor. The good community SIG3, harboring three discriminant species for controls (*Neisseria perflava*, *Rothia mucilaginosa* and *Granulicatella elegans*), and eight of the overall 23 control-related species, it’s the furthest from the bad SIG1, probably collecting different genetic and metabolic pathway features with a potential to counteract COVID-19-related species. Among the 122 species retrieved from DADA2 pipeline (Table S1), 102 were shared among the two cohorts, while twelve and eight were present only in controls and COVID-19 samples, respectively (Figure 1E). Venn diagram relies on the presence of species, not their relative abundance or prevalence, thus, in order to select bacterial species having a plausible and reliable role as biomarkers for COVID-19 and controls, we employed a combination of Volcano plot (Figure 1F), species pairwise comparison (Figure S2, Figure S3), VIP plot and network analysis, resulting in 11 selected species. Bacterial species biomarkers for COVID-19 are *Prevotella salivae*, *Veillonella infantium*, *Prevotella jejuni* and *Soonwooa purpurea* (this latter being present in COVID-19 patients only, Figure 1E). Biomarkers species for healthy oral microbiota are *Neisseria perflava*, *Kallipyga gabonensis*, *Granulicatella elegans*, *Porphyromonas pasteri*, *Gemella taiwanensis*, *Rothia mucilaginosa*, *Streptococcus oralis*.

**Figure 1.**
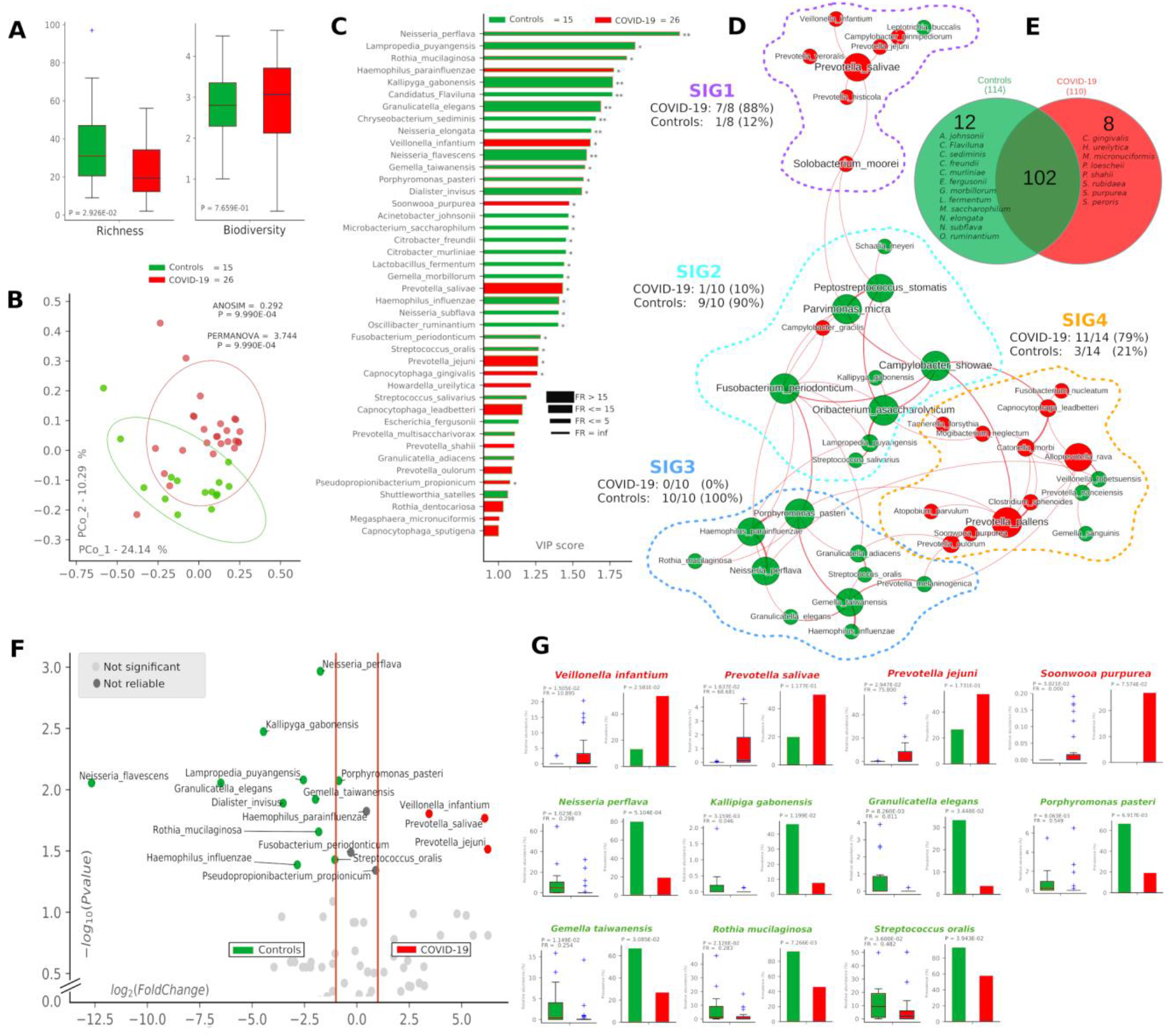
Microbiota composition in COVID-19 and control samples. Alfa- and beta-diversity (panel A and B) of controls (green, n=15) and COVID-19 patients (red, n=26). Variable Importance Plot (VIP, panel C) shows: i) discriminant species after PLS-DA in descending order of VIP score (bar length); ii) the highest relative abundance depending on the cohort (central bar color) and the lowest one (edge bar color); iii) fold ratio (FR) of the highest vs the lowest relative abundance (bar thickness); iv) significant difference after Mann-Whitney U test (non-FDR, * P≤0.05, ** P≤0.01; *** P≤0.001). Network analysis (panel D) shows communities of bacterial species (namely species-interacting groups, SIGs) and their positive (red Pearson coefficient) or negative (blue Pearson coefficient) relative abundances correlation. Nodes are colored according to the cohort harboring the higher relative abundance for a definite species, and node name size is directly proportional to the “keystonness” (importance of a species within the overall network). Edge thickness is inversely proportional to the Pearson P value after 10% Benjamini-Hochberg two-stages FDR, and it is colored according to positive (red) or negative (blue) Pearson coefficient. For each SIG are reported percentages of COVID-19- and controls-related species. Venn diagram (panel E) shows species distribution among the two cohorts considering all of the 122 species (not the core microbiota) retrieved by DADA2 pipeline. Volcano plot (panel F) highlights discriminant oral bacterial species in terms of their fold change (x axis) and cologarithm of Mann-Whitney U test P value (non-FDR) (y axis): species with zero relative abundance were not reported. Pairwise analysis (panel G) of selected 11 species (four for COVID-19 - red, and seven for controls - green) depicts significant differences in terms of relative abundance and prevalence. In each sub-graph are reported the P value (from Mann-Whitney U test) and the fold ratio (FR) among COVID-19 and controls.

### Pro-inflammatory cytokines are distinctive for COVID-19 in both oral and serum samples

After defining oral bacterial species as biomarkers of COVID19, we investigated their possible correlation with proinflammatory cytokines eventually involved in a local “cytokine storm”, as found within patients’ bloodstream recently described in literature. Using a panel of 27 cytokines including chemokines and growth factors, we found that COVID-19 patients were significantly distinguishable from controls using non-supervised methods such as Principal Coordinate Analysis (PCoA, *P*=9.9*10^−4^, Figure 2A) and Hierarchical Classification Analysis (HCA, *P*=0.0046, Figure 2B). Aiming at finding discriminant oral cytokines for COVID-19 status, we employed volcano plot (Figure 2C), VIP plot (Figure 2D), finding out seven COVID-19-related discriminant cytokines (IL-6, IL-5, GCSF, IL-2, TNF-α, GMCSF, INF-γ) while only one (IL-12p70) for controls. IL-6 and IL-12p70 were the most discriminant cytokines for COVID-19 and controls, respectively, as confirmed by their pairwise analysis (Figure 2E). Results from serum cytokines profiling from patients with severe symptomatology and complication, highlighted a superimposable cytokine profile to the oral one of patients at the onset of infection (Table 2), resulting in a significant Pearson positive correlation (Figure 2F). In particular, high levels of oral cytokines involved in early antiviral response mirrored cytokine levels in systemic circulation. As reported in literature for the lung, the expression of human ACE2 and TMPRSS2 in mucosal oral samples were downregulated in infected patients (greater than 100 fold) compared with no infected subjects, and no significant association was found with microbiome composition or cytokines profile (data not shown).

**Figure 2.**
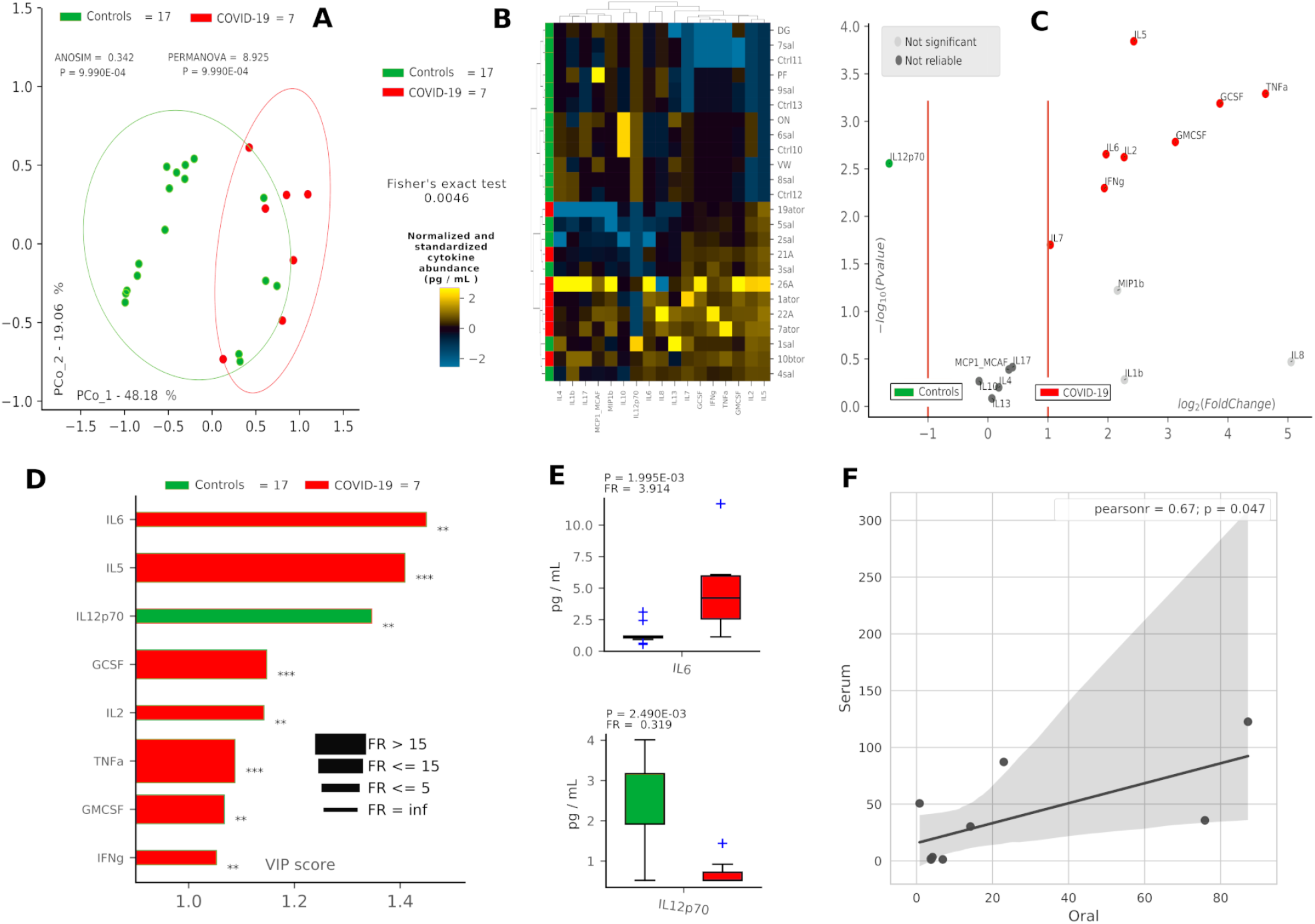
Cytokine pattern in COVID-19 and controls. Cohort separation of controls (green, n=17) and COVID-19 patients (red, n=7) based on cytokine profiles, showed by Principal Coordinate Analysis (PCoA, panel A) and Hierarchical Clusterization Analysis (HCA, panel B), following the Bray-Curtis distance algorithm. Volcano plot (panel C) highlights discriminant oral cytokines in terms of their fold change (x axis) and cologarithm of Mann-Whitney U test P value (non-FDR) (y axis). Variable Importance Plot (VIP, panel D) shows: i) discriminant cytokines after PLS-DA in descending order of VIP score (bar length); ii) the highest cytokine quantity (pg/mL) depending on the cohort (central bar color) and the lowest one (edge bar color); iii) fold ratio (FR) of the highest vs the lowest cytokine quantity (pg/mL) (bar thickness); iv) significant difference after Mann-Whitney U test (non-FDR, * P≤0.05, ** P≤0.01; *** P≤0.001). Pairwise analysis (panel E) of selected two cytokines depicts significant differences in terms of quantity (pg/mL), reporting P value (from Mann-Whitney U test) and fold ratio (FR) among COVID-19 and controls. Pearson linear correlation (panel F) on non-normalized and non-standardized oral (x axis) and serum (y axis) cytokines levels (pg/mL), showing significant positive correlation among the two cytokine patterns. Outlier cytokines having extreme values (IL-1Ra, IL-15, PDGF-bb) were excluded from linear correlation analysis.

**Table 2.**
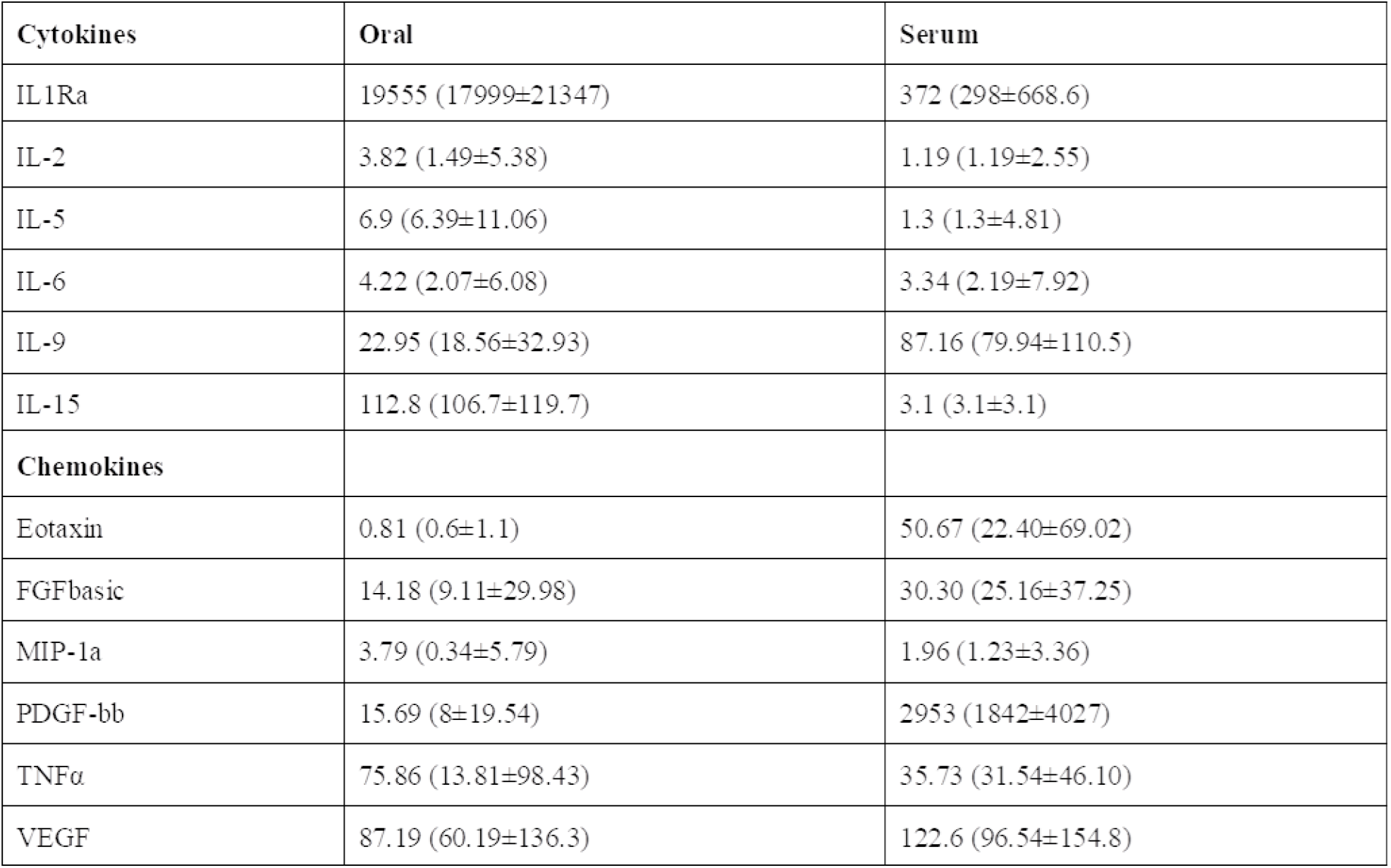
Concentrations (pg/ml) of cytokines and chemokines in patients affected by Covid-19. Immune profile in oral samples from patients at the time of onset and in serum of patients with severe symptomatology. Data are given as medians (quartiles).

| Cytokines | Oral | Serum |
| --- | --- | --- |
| IL1Ra | 19555 (17999±21347) | 372 (298±668.6) |
| IL-2 | 3.82 (1.49±5.38) | 1.19 (1.19±2.55) |
| IL-5 | 6.9 (6.39±11.06) | 1.3 (1.3±4.81) |
| IL-6 | 4.22 (2.07±6.08) | 3.34 (2.19±7.92) |
| IL-9 | 22.95 (18.56±32.93) | 87.16 (79.94±110.5) |
| IL-15 | 112.8 (106.7±119.7) | 3.1 (3.1±3.1) |
| **Chemokines** |  |  |
| Eotaxin | 0.81 (0.6±1.1) | 50.67 (22.40±69.02) |
| FGFbasic | 14.18 (9.11±29.98) | 30.30 (25.16±37.25) |
| MIP-1a | 3.79 (0.34±5.79) | 1.96 (1.23±3.36) |
| PDGF-bb | 15.69 (8±19.54) | 2953 (1842±4027) |
| TNFα | 75.86 (13.81±98.43) | 35.73 (31.54±46.10) |
| VEGF | 87.19 (60.19±136.3) | 122.6 (96.54±154.8) |

### Oral bacterial species topologically counteracts COVID-19-related cytokines

Once found that specific oral cytokines were distinctive for COVID-19, and that the oral cytokine profile was similar to the systemic circulation one, we made a cross-correlation and a network analysis aimed at finding functional clustering and relations among bacterial species and cytokines. Within the network, two distinct communities were formed, separated by a “structural gap” (bunch of negative Pearson correlations) (Figure 3A): the upper community (“GREEN”) harboring 86% species or cytokines from controls, and the lower one (“RED”) hosting 85% species or cytokines having higher abundance in COVID-19 patients (χ^2^= 20.5, *P*<0.00001). Interestingly, keystone species in the GREEN community were *Rothia mucilaginosa* and *Streptococcus oralis*, already evidenced within the good community SIG3 (Figure 1D), while two cytokines (GMCSF and IL-4) were keystone within the RED community. Moreover, within RED cluster we observed a sub-cluster of COVID-19-related species (*Veillonella infantium*, *Prevotella jejuni*, *Streptococcus cristatus*) already seen within the bad community SIG1 (Figure 1D) that were at the farthest distance from GREEN, thus highlighting their functional negative effect. In this proposition, given that unifying oral cytokine and species datasets crunched the overall network structure passing from four to two communities, and that a marked “structural gap” was evidenced among GREEN and RED communities (Figure 3A), the next step was to study all the possible correlations among the single species and the single cytokines, with the intention to highlight oral bacterial species with a potential to counteract COVID-19-related cytokines. A Hierarchical Clusterization Analysys (HCA) correlogram based on Pearson correlation coefficients was performed, resulting in three different clusters of species based on their positive or negative correlation with cytokines (Figure 3B). Cluster1 harbored bad species (such as *Veillonella infantium*, *Streptococcus cristatus*, *Prevotella denticola*, *Atopobium parvulum*) already seen within COVID-19-related communities as SIG1, SIG4 (Figure 1D) and RED (Figure 3A). Cluster2 and Cluster3 contained mostly beneficial species that, conversely, were present within SIG2, SIG3 (Figure 1D) and GREEN (Figure 3A). With this notion in mind, and starting from Pearson coefficients (****r****) of the two cytokines distinctive for COVID-19 (IL-6) and controls (IL-12p70) (Figure 2D), we computed a parameter called C4 (COVID-19 Cytokines Counteracting Coefficient) valuable for each oral bacterial species (Figure 3B):

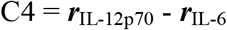

**Figure 3.**
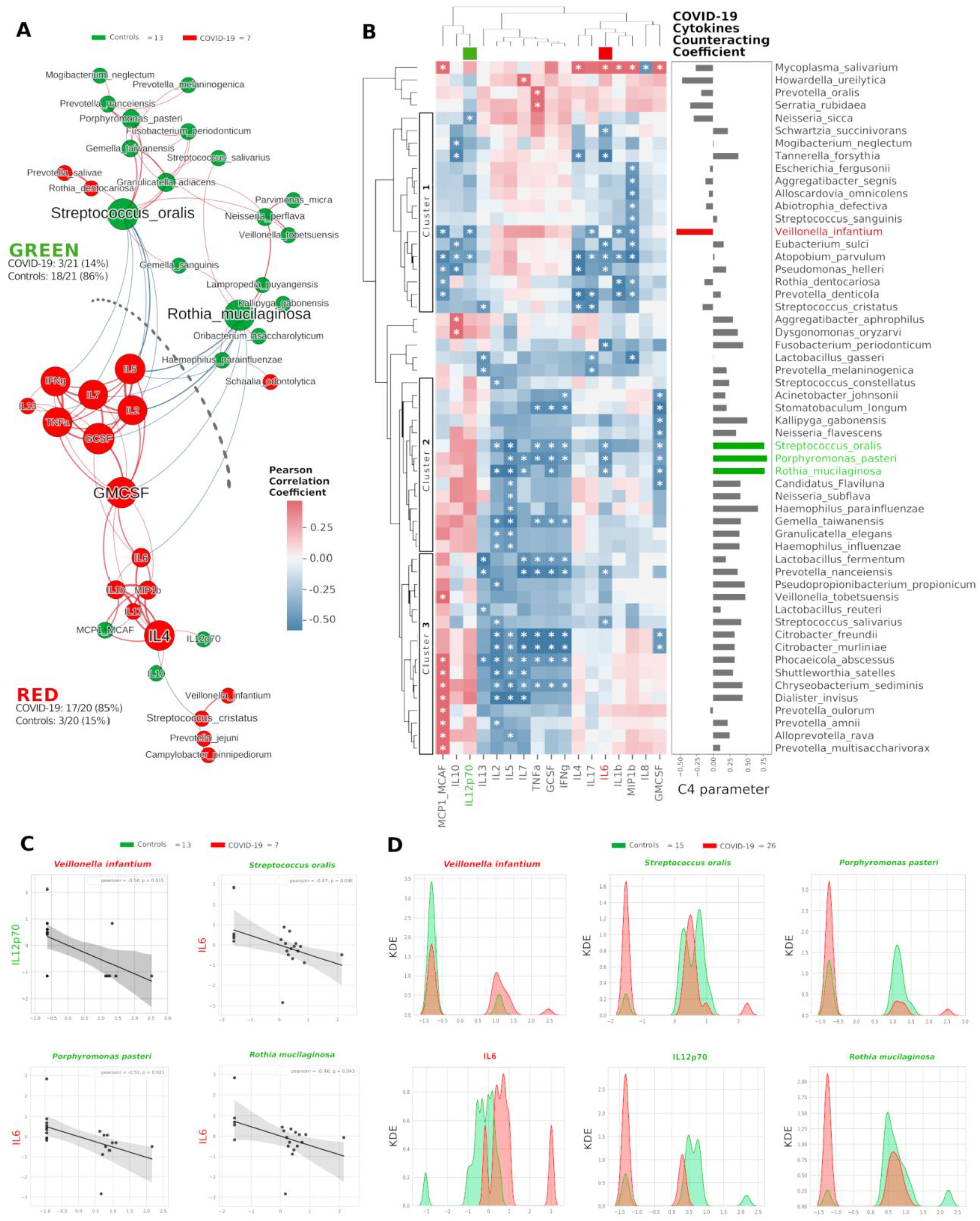
(previous page). Integration of oral species and cytokines datasets. Network analysis (panel A) shows communities (namely GREEN for controls and RED for COVID-19) of bacterial species and cytokines and their positive (red Pearson coefficient) or negative (blue Pearson coefficient) abundances (relative percentage or pg/mL, respectively for species and cytokines) correlation. Nodes are colored according to the cohort harboring the higher abundance for a definite species or cytokine, and node name size is directly proportional to the “keystonness” (importance of a species or cytokine within the overall network). Edge thickness is inversely proportional to the Pearson P value after 10% Benjamini-Hochberg two-stages FDR, and it is colored according to positive (red) or negative (blue) Pearson coefficient. For each community are reported percentages of COVID-19- and controls-related nodes. Dashed line represents a “structural gap” (a bunch of negative Pearson correlation edges) between GREEN and RED communities. Correlogram (panel B) of bacterial species and cytokines shows positive (red) or negative (blue) Pearson correlation on normalized and standardized abundances. Significant correlation is marked with an asterisk inside each square: only species or cytokines having at least one significant correlation were reported. Dendrograms on the x and y axis were generated following Bray-Curtis similarity, evidencing three different clusters for bacterial species (shown here within white boxes). Cytokines chosen to compute the C4 parameter (IL-12p70, IL-6) are highlighted with a colored box in the top dendrogram. The parameter C4 computed for each species is depicted as barplot length at the right of the correlogram, highlighting the bad species (in red) or the good ones (in green). Scatterplots (panel C) among the four selected species and the two cytokines used to compute the C4 parameter: their abundances are negatively related to one another (normalized and standardized data), as reported by Pearson coefficient and P value (95% confidence interval, gray area). Kernel Density Estimation (KDE) plots (panel D) report on x axis the normalized and standardized abundance of the selected species and cytokines and on y axis the subjects’ distribution, divided by COVID-19 (red) and controls (green).

Averaging all C4 values within each cluster of the HCA correlogram, resulted in C4_cluster1_ = −0.017, C4_cluster2_ = 0.472, C4_cluster3_ = 0.301, with a significant difference among Cluster1 and Cluster2 (*t* = - 2.72764, *P*=0.0084, Figure 3B). Noteworthy, the species having the highest C4 values were *Streptococcus oralis*, *Porphyromonas pasteri* and *Rothia mucilaginosa*, species previously found in SIG3 and GREEN communities. The detrimental species *Veillonella infantium*, already found in the RED sub-cluster (Figure 3A) and within SIG1 (Figure 1D), had the highest negative C4 value, thus representing a plausible helper species for COVID-19 onset. In order to confirm the relation among these four species and the involved cytokines, we performed Pearson linear correlation scatterplots (Figure 3C) and Kernel Density Estimation (KDE) area plots (Figure 3D). Linear scatterplots confirmed the expected negative correlation among beneficial species and the pro-inflammatory IL-6, thus meaning that higher amounts of these bacteria could lower the pro-inflammatory oral environment. KDE plots measured patients’ distribution along the abundance of the four selected species or the two cytokines, evidencing how *Rothia mucilaginosa* and *Porphyromonas pasteri*, having a clear superimposition of two peaks centered on the same value for COVID-19 (red) and controls (green), would act differently from *Streptococcus oralis*, which presents two green peaks mutually excludable from the single red one (Figure 3D). The information provided by KDE plots would thus be compulsory for a plausible fine-tuning regulation of species relative abundances in the oral cavity against COVID-19 onset. Taking into consideration the results from Figure 1 and Figure 3, we selected three species as potential counteractors of COVID-19, namely *Streptococcus oralis*, *Rothia mucilaginosa* and *Porphyromonas pasteri*. These species topologically grouped within SIG3, together with other possible candidates considerable as “helpers” to their positive function: *Granulicatella adiacens*, *Granulicatella elegans*, *Gemella taiwanensis*, *Neisseria perflava*. The collective information gathered from network analysis, HCA correlograms, and KDE plots integrating species and cytokines datasets, would thus be amenable for a specific probiotic formulation committed against COVID-19 onset and/or COVID-19-related cytokines.

### Consortia of bacteria and cytokines predict COVID-19 status and its neurological symptoms

After selecting eleven bacterial species (Figure 1) and eight cytokines (Figure 2) as biomarkers for COVID-19 and controls, along with their topological relationships (Figure 3), we focused our attention to their predictive power. Employing the Receiver Operating Characteristic (ROC) metric to evaluate classifier output quality using a 5-fold cross-validation, we evidenced how single cytokines gave a higher power when unified to single species in predicting COVID-19 status (Figure 4A, Figure 4B), showing a significant higher averaged Area Under Curve (AUC) (AVG_AUCavg(Cytokines_Species)_ = 0.891; AVG_AUCavg(Species)_ = 0.637; Mann-Whitney U test, two-tailed *P*=0.012). Moreover, the first five ROC curves in Figure 2B, representing the ROCs having the best AUC values, were devoted to only cytokines, thus confirming their importance in defining the disease status better than species. Aiming at finding the best consortium of bacterial species (n=11) or species and cytokines (n=19) able to predict the COVID-19 status, a combinatorial calculation was performed (supplementary Excel File). A total of 2047 and 524287 combinations were retrieved for species (Figure 2C) and species plus cytokines (Figure 2D), respectively, and also in this scenario the best averaged AUC value was significantly higher (Mann-Whitney U test, two-tailed *P*=0.012) for “consortia’’ of species plus cytokines (AVG_AUCavg(Cytokines_Species)_ = 0.995), other than considering proper consortia formed by species alone (AVG_AUCavg(Cytokines_Species)_ = 0.932). Interestingly, majority of consortia showing ROC curves with the highest values of AUC, specificity and sensitivity (namely 0.995, 0.990 and 1.000, Figure 4D), were those encompassing a balanced ratio of species and cytokines that were beneficial (*Prevotella salivae*, *Streptococcus oralis*, *Rothia mucilaginosa*, *Gemella taiwanensis*, *Kallipyga gabonensis*, *Granulicatella elegans*, IL-12p70) or detrimental (*Prevotella jejuni*, *Soonwooa purpurea*, *Veillonella infantium*, TNF-α, INF-γ, IL-2, IL-6, IL-5, GCSF, GMCSF). After assessing that definite combinations of our selected bacterial species, alone or added to oral cytokines, were able to discriminate COVID-19 status, we aimed at parametrizing each single subject for a general COVID-19 susceptibility. To this aim, we created two parameters, named BacCOV and CytoCOV, based on the abundances of selected bacterial species or cytokines (shown in Figure 1G, Figure 2D). More precisely, in order to have a bidimensional representation for both cohorts, usable to visually predict the propensity of a putative unknown subject to COVID-19, we computed for each subject: i) the BacCOV_GREEN and CytoCOV_GREEN averaging the abundances of beneficial species (n=7) or species plus cytokines (n=7+1); ii) the BacCOV_RED and CytoCOV_RED value averaging the abundances of detrimental species (n=4) or species plus cytokines (n=4+7) (Table S2, Table S3). These four values were used to generate L-shaped graphs, in which we set x-axis and y-axis thresholds (computed as in Table S4) in order to ease a clinical usage for subjects’ COVID-19 susceptibility (Figure 4E, Figure 4F). Even if BacCOV significantly divided the two cohorts (*P* = 2.1*10^−3^), a one-order higher significant separation was obtained with CytoCOV (*P* = 4*10^−4^), correctly classifying 71% of COVID-19 patients (Figure 4F). Thus, the CytoCOV parameter would be easily employed in clinics to assess COVID-19 susceptibility for an unknown subject, through the following passages: i) assaying the abundances of 11 bacterial species and 8 cytokines; ii) computing the CytoCOV parameter (Table S3); iii) using the boundaries provided in Table 3.

**Figure 4.**
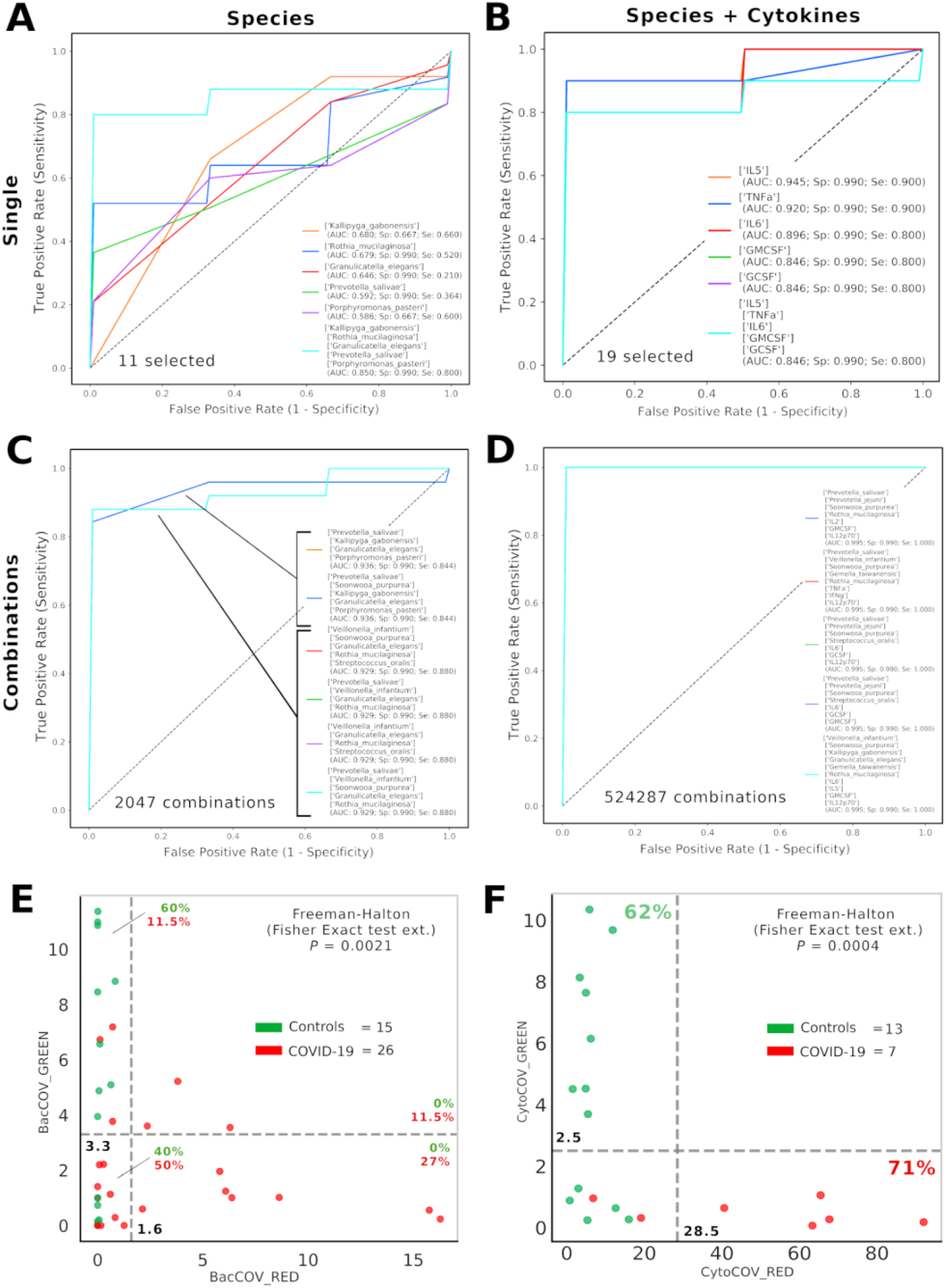
Consortia of bacteria and cytokines predict COVID-19 status. Receiver Operating Characteristic (ROC) with 5-fold cross validation and C-Support Vector Machine classifier was employed to assess the best oral bacterial species (among 11 selected, panel A) or species plus cytokines (among 19 selected, panel B) able to predict COVID-19 status. Each panel reports the best five Area Under Curve (AUC) values in descending order (see the inset legend also showing specificity, Sp, and sensitivity, Se, for each ROC curve), plus a sixth curve encompassing the preceding five grouped. Combinations were computed for selected species (n=11, 2047 combinations, panel C) and species plus cytokines (n=19, 524287 combinations, panel D), and the best “consortia’’ predicting COVID-19 are shown along with their AUC, Sp and Se values. BacCOV (panel E) and CytoCOV (panel F) parameters were computed and divided into their GREEN (species and cytokines higher in controls) and RED (species and cytokines higher in COVID-19) components, and employed to generate scatterplot 2D graphs. Abundance thresholds (computed as in Table S4) are shown as dotted grey lines, and their values reported in bold. In each quadrant of the panels E and F are reported the percentages of controls (green) or COVID-19 (red) subjects.

**Table 3.** Boundaries for CytoCOV parameter, as evidenced in Figure 4F, in order to assess subjects’ COVID-19 predicted susceptibility. Percentage values represent subjects’ fraction.

| | CytoCOV_RED $\geq 28.5$<br>CytoCOV_GREEN $< 2.5$ | CytoCOV_RED $< 28.5$<br>CytoCOV_GREEN $\geq 2.5$ | CytoCOV_RED $< 28.5$<br>CytoCOV_GREEN $< 2.5$ |
| --- | --- | --- | --- |
| Controls | 0 | 62% | 38% |
| COVID-19 | 71% | 0 | 29% |

Thirty-five percent of our COVID-19 patients (9/26) presented, as comorbidity, a neurological involvement like ageusia or hyposmia. Applying the same analysis used previously, even if these patients did not possess a distinctive oral microbiota composition (data not shown), significant higher levels of detrimental species such as *Prevotella jejuni* and *Streptococcus cristatus* were found (Figure 5A). Interestingly, these species were found within the RED community in the combined species/cytokines network (Figure 3A). Averaged Area Under Curve (AUC) values of ROC curves regarding these two selected species were 0.864 and 0.775, respectively (Figure 5B), while their combination ensured an accurate prediction of 67% of patients with neurological symptoms (confusion matrix, Figure 5C). Interestingly, these two species were positively and significantly related (*r* = 0.35, *P*=0.027, Figure 5D), so we employed a bidimensional representation of their relative abundances in order to assess patients’ neurological symptoms susceptibility (Figure 5E). An unknown COVID-19 patient would thus be susceptible of neurological symptoms if harboring *Prevotella jejuni* and *Streptococcus cristatus* at relative abundances higher than or equal to 17.1% or 14.4%, respectively (thresholds computed as in Table S5, Fisher Exact test *P* = 1.9*10^−3^).

**Figure 5.**
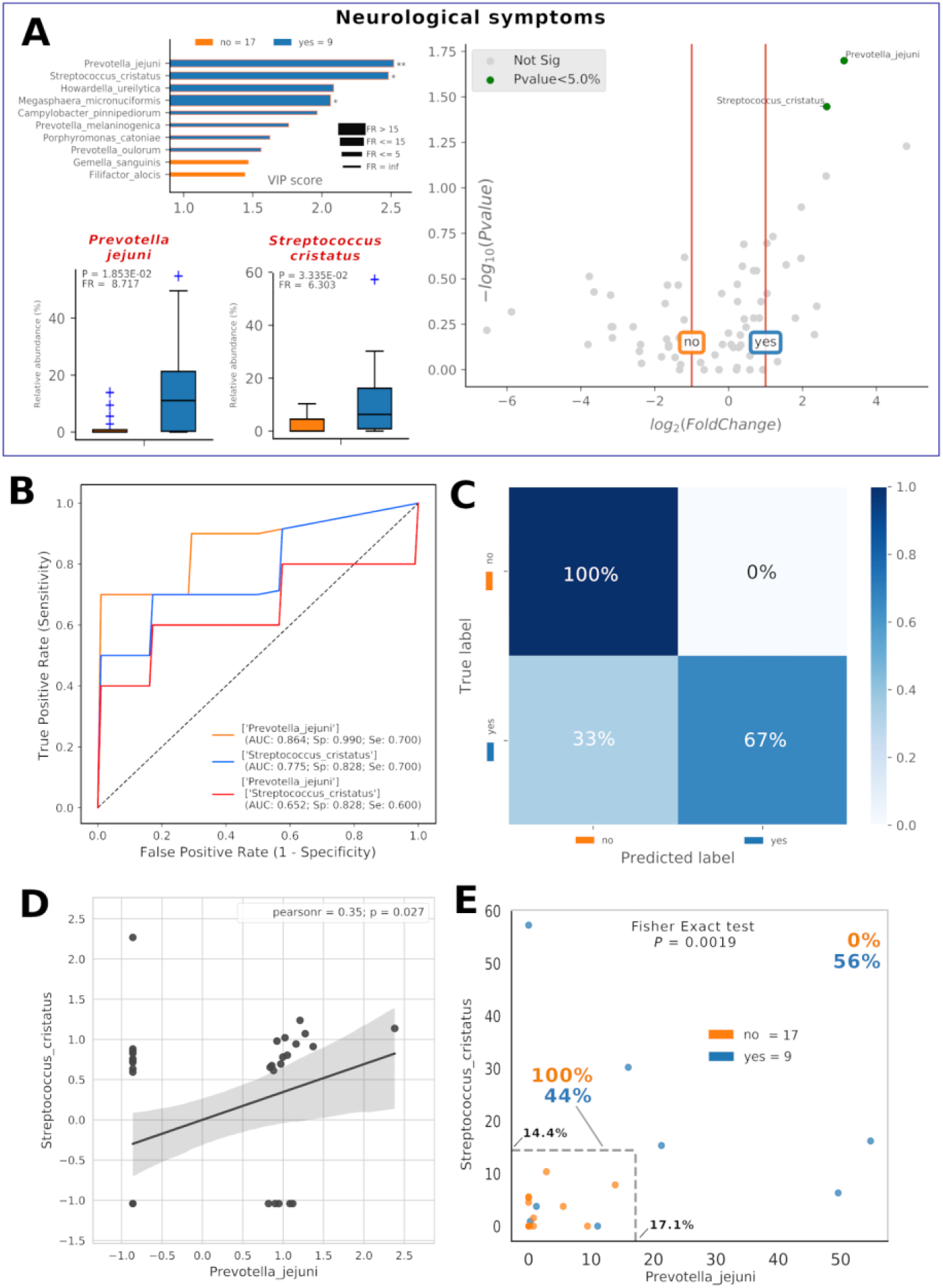
Consortia of bacteria predict COVID-19 neurological symptoms. Mixed graphs for COVID-19 patients having neurological symptoms (panel A) are made of VIP plot, volcano plot, and boxplot of relative abundances for selected species. Description for these plots as in Figure 1 and Figure 2 captions. Receiver Operating Characteristic (ROC) with 5-fold cross validation and C-Support Vector Machine classifier (panel B) was built for each one of the selected species plus their combined effect on predicting the presence of neurological symptoms (“yes”, blue) in respect to their absence (“no”, orange). Panel B reports Area Under Curve (AUC), specificity (Sp), and sensitivity (Se) values for each ROC curve. Confusion matrix (panel C) was employed to evaluate the quality of the output of the C-Support Vector Machine classifier used to generate ROC curves. The diagonal elements represent the percentage of patients (referred to the blue shadowed sidebar) for which the predicted label (x axis) is equal to the true label (y axis), while off-diagonal elements are those that are mislabeled by the classifier. Scatterplot (panel D) among the two selected species shows how their relative abundances are positively related to one another (normalized and standardized data), as reported by Pearson coefficient and P value (95% confidence interval, gray area). Raw relative abundances of selected species were used to generate a scatterplot (panel E) to ease a clinical interpretation of patients’ distribution into the 2D space. Abundance thresholds (computed as in Table S5) are shown as truncated dotted grey lines, and their values reported in bold: percentages of patients separated by these boundaries are reported in bold.

## DISCUSSION

COVID-19 proved to be an important threat to our lives, harnessing more than 60M cases worldwide and around 1.5M global deaths, as for November 2020 ^29^. The major issue when coping with the COVID-19 etiological agent, SARS-CoV-2 betacoronavirus, resides in its high spreading ability, which is mediated by several interpersonal factors, such as oral droplets ^30,31^. In this proposition, the oral cavity should be considered a preferential route for SARS-CoV-2 entry or transmission ^32,30^, especially in nosocomial environments at risk ^33^, with potential involvement for extrapulmonary sites (such as the brain ^5^ and gastrointestinal tract) ^32,34.^ Here we investigated the oral microbiota composition and cytokines in COVID-19 subjects and matched controls. As for November 2020, only three experimental studies focused on the COVID-19-related intestinal bacterial microbiota ^35,36^, and on fungal one ^37^, leaving unanswered questions on the oral district. The new approach of social network analysis allowed us to properly merge species and cytokines datasets, providing definite bacterial consortia as biomarkers for COVID-19 status or its related neurological symptoms, and providing also new parameters for theranostics purposes against COVID-19. Our results especially suggest how a minimal consortium of beneficial species (*Prevotella salivae*, *Streptococcus oralis*, *Rothia mucilaginosa*, *Gemella taiwanensis*, *Kallipyga gabonensis*, *Granulicatella elegans*) could be used orally as local probiotics to counteract COVID-19 symptoms and cytokine storm, which is typical in COVID-19 patients ^38,4.^ In a recent study genera *Rothia*, *Streptococcus* and *Veillonella* were positively related to COVID-19 in feces ^35^, while here we found how some oral species belonging to these genera exert a counteracting effect on COVID-19 cytokine storm ^4^. This discrepancy is noteworthy, because future studies dealing with different body districts should consider shotgun sequencing or 16S targeted sequencing to reach the species level, assuring a proper functional description (e.g., combination of microbiota and cytokine data) for clinical use. Being attractive for the ease of sampling, as demonstrated in international projects such as HMP, the oral swab sampling (touching tongue, palatum and cheeks) would be a valid alternative to the currently used specimens (e.g. feces, blood) to assess the microbiota compositional differences in a disease, giving reliable insights on a subject susceptibility. Even if with intrinsic limitations due to the number of subjects involved, and the missing point of a shotgun implementation to ascertain gene and/or pathways differences among controls and COVID-19, our study would give a hint to the significance of the oral microbiota restoration during pandemic as a public health intervention ^31,39–42^.

## REFERENCES

1. Li, J. et al. Clinical features of familial clustering in patients infected with 2019 novel coronavirus in Wuhan, China. Virus Res. 286, 198043 (2020).

2. Srivastava, P. & Gupta, N. Clinical Manifestations of Corona Virus Disease. Clinical Synopsis of COVID-19 31–49 (2020) doi:10.1007/978-981-15-8681-1_3.

3. de la Rica, R., Borges, M. & Gonzalez-Freire, M. COVID-19: In the Eye of the Cytokine Storm. Front. Immunol. 11, 558898 (2020).

4. Jose, R. J. & Manuel, A. COVID-19 cytokine storm: the interplay between inflammation and coagulation. Lancet Respir Med 8, e46–e47 (2020).

5. Gupta, A. et al. Extrapulmonary manifestations of COVID-19. Nat. Med. 26, 1017–1032 (2020).

6. Wang, D. et al. Clinical Characteristics of 138 Hospitalized Patients With 2019 Novel Coronavirus-Infected Pneumonia in Wuhan, China. JAMA 323, 1061–1069 (2020).

7. Chen, X. et al. Detectable serum SARS-CoV-2 viral load (RNAaemia) is closely associated with drastically elevated interleukin 6 (IL-6) level in critically ill COVID-19 patients. doi:10.1101/2020.02.29.20029520.

8. Qin, C. et al. Dysregulation of Immune Response in Patients with COVID-19 in Wuhan, China. SSRN Electronic Journal doi:10.2139/ssrn.3541136.

9. Braz-Silva, P. H., Pallos, D., Giannecchini, S. & To, K. K. SARS-CoV-2: What can saliva tell us? Oral Diseases (2020) doi:10.1111/odi.13365.

10. Leung, E. C., Chow, V. C., Lee, M. K. & Lai, R. W. Deep throat saliva as an alternative diagnostic specimen type for the detection of SARS-CoV-2. Journal of Medical Virology (2020) doi:10.1002/jmv.26258.

11. To, K. K.-W. et al. Temporal profiles of viral load in posterior oropharyngeal saliva samples and serum antibody responses during infection by SARS-CoV-2: an observational cohort study. Lancet Infect. Dis. 20, 565–574 (2020).

12. To, K. K.-W. et al. Consistent Detection of 2019 Novel Coronavirus in Saliva. Clin. Infect. Dis. 71, 841–843 (2020).

13. Herrera, D., Serrano, J., Roldán, S. & Sanz, M. Is the oral cavity relevant in SARS-CoV-2 pandemic? Clin. Oral Investig. 24, 2925–2930 (2020).

14. Feng, Z., Wang, Y. & Qi, W. The Small Intestine, an Underestimated Site of SARS-CoV-2 Infection: From Red Queen Effect to Probiotics. doi:10.20944/preprints202003.0161.v1.

15. Zanotta, N. et al. Up-regulation of the monocyte chemotactic protein-3 in sera from bone marrow transplanted children with torquetenovirus infection. J. Clin. Virol. 63, 6–11 (2015).

16. Ma, D. et al. Expression of SARS-CoV-2 receptor ACE2 and TMPRSS2 in human primary conjunctival and pterygium cell lines and in mouse cornea. Eye 34, 1212–1219 (2020).

17. Campisciano, G. et al. Shifts of subgingival bacterial population after nonsurgical and pharmacological therapy of localized aggressive periodontitis, followed for 1 year by Ion Torrent PGM platform. Eur. J. Dent. 11, 126–129 (2017).

18. Derosa, L. et al. Gut Bacteria Composition Drives Primary Resistance to Cancer Immunotherapy in Renal Cell Carcinoma Patients. Eur. Urol. 78, 195–206 (2020).

19. Li, M. et al. Symbiotic gut microbes modulate human metabolic phenotypes. Proc. Natl. Acad. Sci. U. S. A. 105, 2117–2122 (2008).

20. Merico, D., Gfeller, D. & Bader, G. D. How to visually interpret biological data using networks. Nat. Biotechnol. 27, 921–924 (2009).

21. Berry, D. & Widder, S. Deciphering microbial interactions and detecting keystone species with co-occurrence networks. Front. Microbiol. 5, 219 (2014).

22. Faust, K. et al. Microbial co-occurrence relationships in the human microbiome. PLoS Comput. Biol. 8, e1002606 (2012).

23. Lozupone, C. A., Stombaugh, J. I., Gordon, J. I., Jansson, J. K. & Knight, R. Diversity, stability and resilience of the human gut microbiota. Nature 489, 220–230 (2012).

24. Faust, K. & Raes, J. Microbial interactions: from networks to models. Nat. Rev. Microbiol. 10, 538–550 (2012).

25. Iebba, V. et al. Combining amplicon sequencing and metabolomics in cirrhotic patients highlights distinctive microbiota features involved in bacterial translocation, systemic inflammation and hepatic encephalopathy. Sci. Rep. 8, 8210 (2018).

26. Blondel, V. D., Guillaume, J.-L., Lambiotte, R. & Lefebvre, E. Fast unfolding of communities in large networks. Journal of Statistical Mechanics: Theory and Experiment vol. 2008 P10008 (2008).

27. Lambiotte, R., Delvenne, J.-C. & Barahona, M. Random Walks, Markov Processes and the Multiscale Modular Organization of Complex Networks. IEEE Transactions on Network Science and Engineering vol. 1 76–90 (2014).

28. Anderson, M. J. & Walsh, D. C. I. PERMANOVA, ANOSIM, and the Mantel test in the face of heterogeneous dispersions: What null hypothesis are you testing? Ecological Monographs vol. 83 557–574 (2013).

29. Dong, E., Du, H. & Gardner, L. An interactive web-based dashboard to track COVID-19 in real time. Lancet Infect. Dis. 20, 533–534 (2020).

30. Netz, R. R. & Eaton, W. A. Physics of virus transmission by speaking droplets. Proc. Natl. Acad. Sci. U. S. A. 117, 25209–25211 (2020).

31. Bao, L. et al. Oral Microbiome and SARS-CoV-2: Beware of Lung Co-infection. Front. Microbiol. 11, 1840 (2020).

32. Jiao, L. et al. The gastrointestinal tract is an alternative route for SARS-CoV-2 infection in a nonhuman primate model. Gastroenterology (2020) doi:10.1053/j.gastro.2020.12.001.

33. Kumbargere Nagraj, S. et al. Interventions to reduce contaminated aerosols produced during dental procedures for preventing infectious diseases. Cochrane Database Syst. Rev. 10, CD013686 (2020).

34. Trottein, F. & Sokol, H. Potential Causes and Consequences of Gastrointestinal Disorders during a SARS-CoV-2 Infection. Cell Reports vol. 32 107915 (2020).

35. Gu, S. et al. Alterations of the Gut Microbiota in Patients With Coronavirus Disease 2019 or H1N1 Influenza. Clinical Infectious Diseases (2020) doi:10.1093/cid/ciaa709.

36. Zuo, T. et al. Alterations in Gut Microbiota of Patients With COVID-19 During Time of Hospitalization. Gastroenterology vol. 159 944–955.e8 (2020).

37. Zuo, T. et al. Alterations in Fecal Fungal Microbiome of Patients With COVID-19 During Time of Hospitalization until Discharge. Gastroenterology vol. 159 1302–1310.e5 (2020).

38. Guo, Y.-R. et al. The origin, transmission and clinical therapies on coronavirus disease 2019 (COVID-19) outbreak - an update on the status. Mil Med Res 7, 11 (2020).

39. Patel, J. & Sampson, V. The role of oral bacteria in COVID-19. Lancet Microbe 1, e105 (2020).

40. Chen, X. et al. The microbial coinfection in COVID-19. Appl. Microbiol. Biotechnol. 104, 7777–7785 (2020).

41. Sampson, V., Kamona, N. & Sampson, A. Could there be a link between oral hygiene and the severity of SARS-CoV-2 infections? British dental journal vol. 228 971–975 (2020).

42. Al-Khatib, A. Oral manifestations in COVID-19 patients. Oral Diseases (2020) doi:10.1111/odi.13477.

